# A mechanistic model of early centrosome separation during prophase

**DOI:** 10.1101/2025.09.03.673972

**Authors:** Subhendu Som

## Abstract

Initial centrosome separation during prophase is crucial for faithfully completing the mammalian mitosis. Though kinesin-5 pushes the two centrosomes apart in prometaphase by sliding the inter-centrosomal anti-parallel microtubules, it is still under debate that whether kinesin-5 plays any role in the prophase centrosome separation. It is also controversial that whether dyneins separate the centrosomes by pulling the centrosomal microtubules from two opposite directions. Therefore, the active drivers involved in the early centrosome separation are still elusive. Interactions of the centrosomal microtubules with cellular objects (e.g. cortex, cell wall and nucleus) generate forces on the centrosome, thereby promotes centrosomal movements. Using an *in-silico* model, we show that a concerted interplay of centrosomal microtubules with cortex, cell wall and nucleus can promote centrosome separation during prophase.

## 1. Introduction

A prerequisite for bipolar spindle assembly is the optimal separation of mother and daughter centrosomes (CSs) around the nucleus [1]. In the mammalian cells, centrosome separation occurs via two steps: initially, an angular separation (with respect to the center of the nucleus) is established between the centrosomes during prophase [1, 2]. Subsequently, in the beginning of prometaphase, the nuclear membrane disappears and the arrays of anti-parallel microtubules (MTs) from the two centrosomes emerge. Bipolar Kinesin-5, crosslink and walks along the anti-parallel microtubules, thereby pushing them away from each other [1, 2]. Though kinesin-5 driven late stage separation of the centrosomes is well regarded in the scientific community, controversy remains whether there is a role of kinesin-5 in the establishment of the centrosomal separation during prophase [3, 4]. If kinesin-5 is crucial, it would engage the antiparallel arrays of microtubules between the centrosomes leading to the separation. Nevertheless, no such dependency is found in the experiments [4]. In addition, existing literature argues that kinesin-5 is nonessential for centrosome separation during prophase in *C. elegans and Drosophila embryos* [5, 6, 7]. Thus, kinesin-5 may not be an active driver for establishing the early centrosome separation.

To elucidate the centrosome separation further, several experiments were carried out testing the function of dynein as the leading force generator [8, 9, 10, 11, 12, 13]. Dyneins residing along the cell cortex or on the outer nuclear membrane engage with the centrosomal microtubules, thereby pull on the centrosome with force ~1-5 pN [14, 15, 16]. In the event centrosomes are pulled in the opposite directions, the action serves as a plausible mechanism for separating the centrosomes. Yet, the outcomes of the previous research are puzzling, as some reports established the positive role of dynein in centrosome separation [10, 11, 12] contradicting the neutral role of dynein reported elsewhere [8, 9]. Additionally, experiments on lower eukaryotes (e.g. yeast) suggest that dynein plays no role determining the centrosomal separation during prophase [17, 18]. Naturally, question arises whether dyneins participate in the centrosome separation at all? Investigations were also carried out to understand the role of actin in the centrosome separation, but the available literature suggests that actin can act as a passive driver only, in the entire separation process [19, 20, 21].

To shed light on the centrosome separation, we perform an *in-silico* study considering three-dimensional model resembling the mammalian cell. Throughout this study, kinesin-5 driven sliding of antiparallel microtubules from opposite centrosome is not taken into account. Upon implementing all major interactions of the microtubules with the cell-cortex and nucleus, we find that centrosomes move independently and separate by ~85^0^. The separation increases significantly (~85^0^ to ~140^0^) when the microtubule-centrosome interactions are incorporated. Microtubule-centrosome interactions generate repulsive forces between the two centrosomes as the microtubules from one centrosome hit the other [22, 23, 24]. Simulation also shows that if the microtubule-dynein interactions, found in cortex or on nuclear envelope, are removed completely, the results remain unchanged. Therefore, this article summarizes the pathway through which the initial centrosome separation can be achieved without the help of kinesin-5 and dynein.

## 2. Model and Simulation

In order to model the cell, nucleus, centrosome, microtubule, cortex and the microtubule forces, the exact same mechanisms are followed as they are in our earlier article [25]. Although we describe the modelings here once more, for details see the literature [25]. We consider cell, nucleus and centrosome as 3D spheres having radii *r*_*cell*_, *r*_*nuc*_ and *r*_*cs*_ respectively (see table 1). Nucleus is static and kept at the cell-center throughout the simulation. Steric repulsion is introduced between centrosome-centrosome, centrosome-nucleus and centrosome-cell wall. At the onset of the simulation, two centrosomes are in touch with each other and localized in the perinuclear region (close to nucleus). Microtubules are semi-flexible polymers and nucleated from the two centrosomes uniformly in all directions. Here we consider 100 microtubules per centrosome. The dynamic instability of the microtubules is implemented using four characteristic parameters i.e. *v*_*g*_ (polymerization speed), *v*_*s*_ (depolymerization speed), *f*_*c*_ (catastrophe rate) and *f*_*r*_ (rescue rate) [26]. *f*_*c*_ promotes catastrophe of the growing microtubules and *f*_*r*_ promotes re-growth of the shrinking microtubules. Microtubules interact with the various cellular objects and thereby apply forces on the microtubule organizing centers (here the centrosomes) as shown in fig. 1. Cell cortex applies an elastic pushing force on the microtubule-tips [27, 28]. If *K*_*cor*_ and 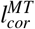 are the spring constant of the cortex and the penetration length of a microtubule in the cortex, respectively, the force arising from the deformation of the cortex is given by 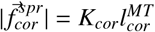. Cortical dyneins bind and walk toward the −ve end of the microtubules, thereby generating pulling forces on the centrosomes [29, 30]. If 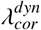 and 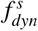 are the linear density of dynein along the microtubule and the force produced by a single dynein, respectively, the pulling force would be 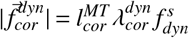. Cell wall appears as barrier to the growing microtubules, hence producing pushing forces 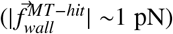 on the microtubule-tips [31]. A resistance to the growing microtubule from the cell wall can either buckle the microtubule or let it slide along the cell-periphery [24, 32, 33]. Although, angle between the microtubule and the cell wall can favor one of the possibilities, an equal probability for buckling and sliding is assumed for simplicity [25]. A bent microtubule, may apply first order Euler buckling force(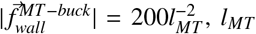 is the total microtubule length) on the centrosome [34, 35, 14] directed along the line joining the microtubule-tip and the centrosome. Cortical dyneins reel in the sliding microtubule, producing pulling force (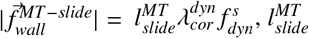 is the length of the sliding portion of the microtubule) on the centrosome [14, 29, 30] directed along the line joining the centrosome and the point of entry of the microtubule in the cortex. Since nucleus is static, nuclear envelope (NE) also acts as a barrier to the growing microtubules [36, 37], thereby producing 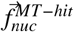 and 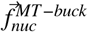 on the microtubule-tips in the similar manner as 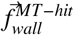 and 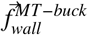, respectively. Like-wise, dyneins residing on the NE [16, 11] produce sliding force 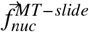 on the microtubule-tips.

**Table 1:**
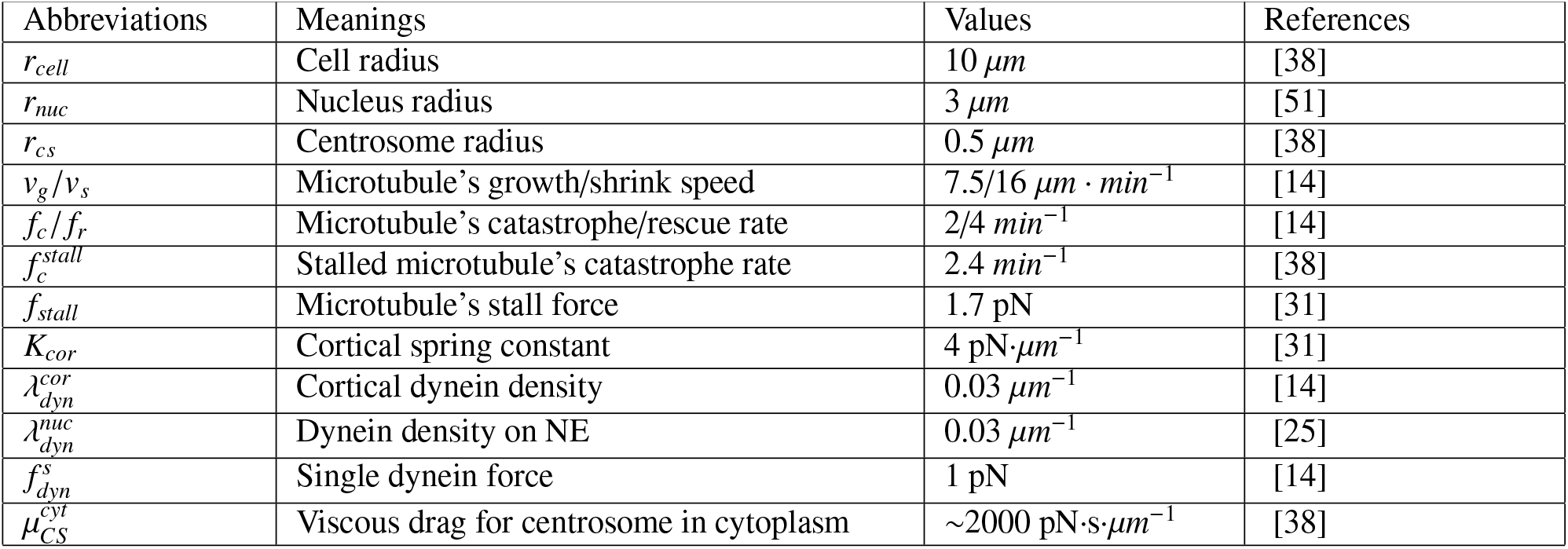
Model Parameters.

**Figure 1:**
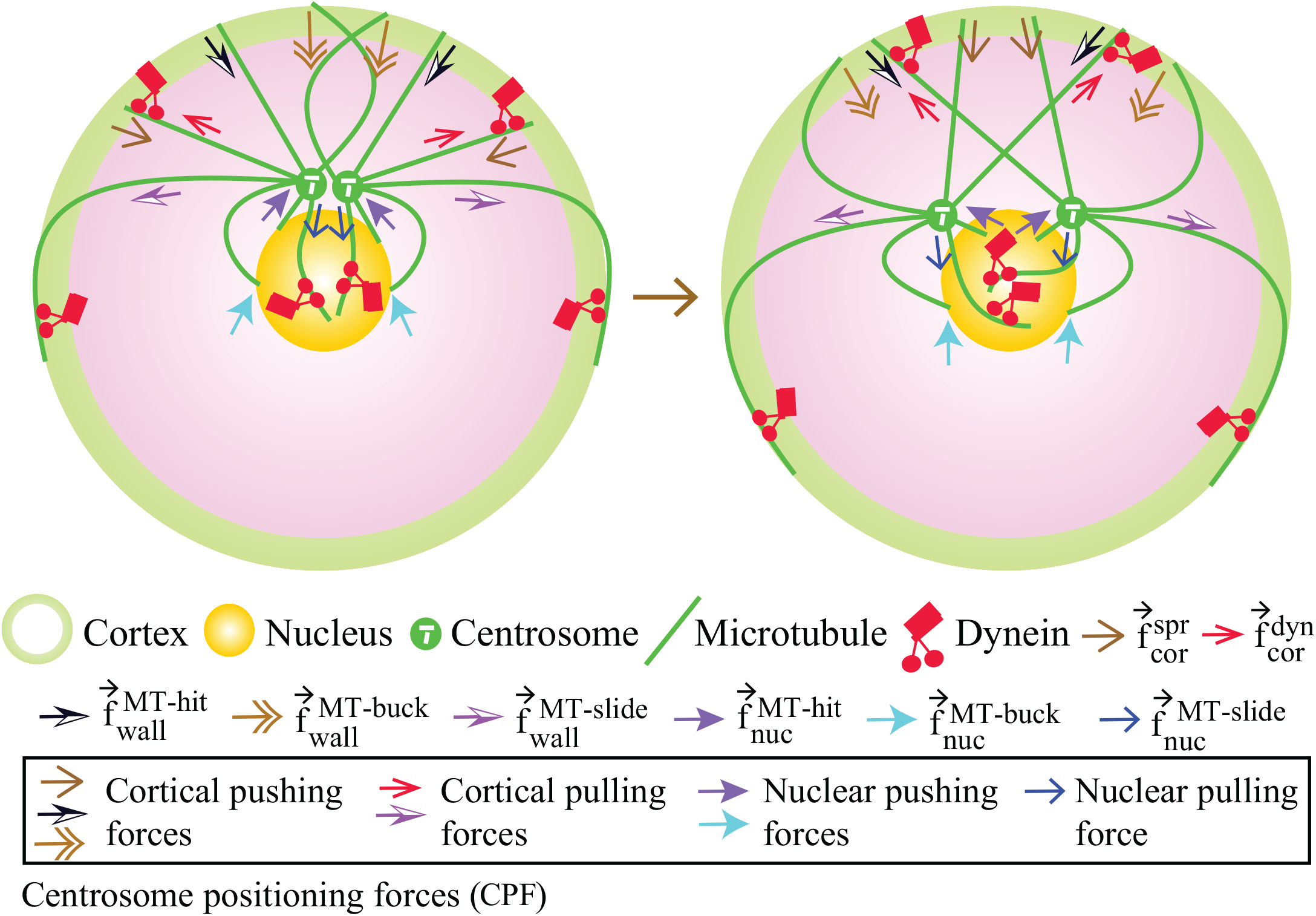
Schematic showing various molecular forces acting on the centrosomes. These forces restrict the centrosomal movements within the perinuclear region.

When loads via 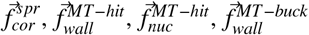 and 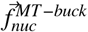 are applied on the microtubule-tips, polymerization speed *v*_*g*_ and catastrophe rate *f*_*c*_ of the microtubule change in the following manner [38]: *v*_*g*_ = *v*_*g*0_ exp(− *f*_*load*_/*f*_*stall*_), where, *f*_*load*_ is the magnitude of the load, *f*_*stall*_ is the load that stalls the growth of a single microtubule and *v*_*g*0_ is the unconstrained polymerization speed under zero load. 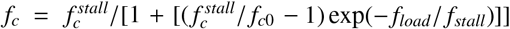, where 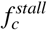 is the rate of catastrophe of a stalled microtubule and 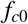 is the catastrophe rate of a free microtubule.

If 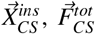 and 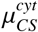 are the instantaneous position of the centrosome, net force acting on the centrosome and viscous drag on the centrosome due to the cytoplasm, respectively, the corresponding equation of motion following Stokes’ law is

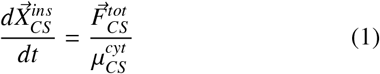

Note that 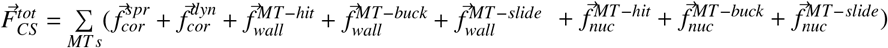.Choosing the time steps 0.01 s (typically much smaller than the time needed to grow a microtubule about the size of the cell), we simulate the system for ~5h which is sufficiently large for obtaining a steady state configuration.

## 3. Results

In our earlier study [25], we showed that the combined effect of all the microtubule driven forces (constituents of 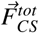) stabilizes the centrosome in the perinuclear region. The stability is governed by a zero-balance of the net force applied on the centrosome along the radial direction in a spherical cell. Forces responsible for proper centrosome positioning, hereafter, referred as CPF (i.e. centrosome positioning forces). In this work, we quantify the role of CPF for centrosome separation. In order to separate the centrosomes around nucleus, CPF must have a component on the plane 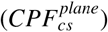 perpendicular to the radius vector 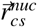 of the centrosome measured from cell center (fig. 2a). Inset of fig. 2a, shows that this component of force remains nonzero with time, while the component of CPF facilitating the radial movement of the centrosome vanishes. However, due to stochastic fluctuation, direction of the 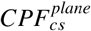 applied on the centrosomes changes frequently and promotes random movement of the centrosomes around the nuclear periphery (fig. 2b). Therefore, when mean angular separation between the centrosomes is measured with respect to the center of the nucleus, it is found to be nearly half (~85^0^) of the complete separation (180^0^) (fig. 2c). Linear separation (distance between the centers of the two centrosomes), plotted in the same figure, also shows a similar characteristics. These outcomes suggest that CPF can facilitate centrosome separation during prophase via independent movement of each centrosome [4]. Since the dynein mediated forces (viz. 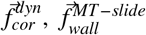 and 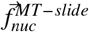) are included in the CPF, it is essential to identify the contributions of these forces in the centrosome separation. All results of fig. 2 remain unaltered (therefore not shown) upon deletion of these forces from the CPF, thereby supporting the neutral role of dynein in the centrosome separation [8, 9]. Quantitatively dynein forces are very small 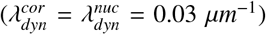,thus suppressed by the other components of the CPF. Hereafter, we remove dyneins from the system and refer CPF as CPF(−).

**Figure 2:**
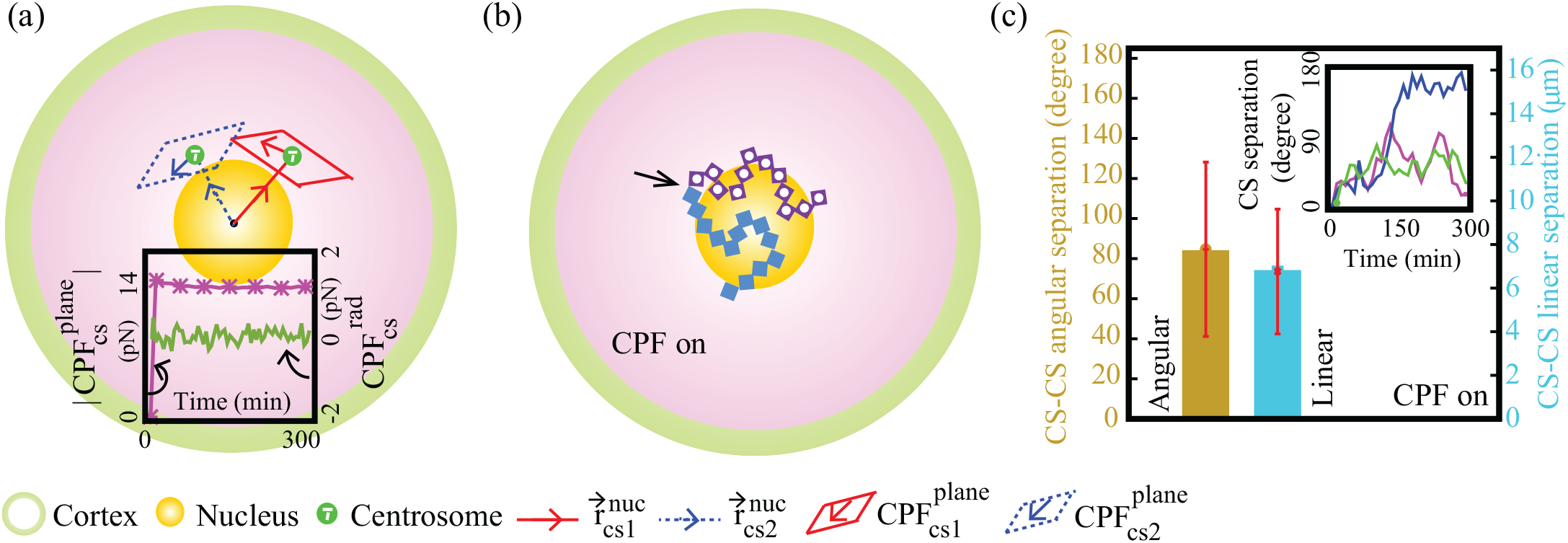
Centrosome positioning forces (CPF) promote the angular movements of the centrosomes around the nucleus. (a) Diagram shows component of CPF 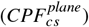 on the plane perpendicular to the radius vector of the centrosome 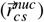.Here CS1 and CS2 denote two different centrosomes. (Inset:) Magnitude of 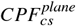 remains almost constant with time, whereas 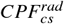,component of CPF along 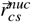 vanishes. (b) Centrosomes move independently (square and dot-square) under the effect of CPF. Arrow represents the starting point of the two centrosomes. (c) Quantification of centrosome separation driven by CPF. (Inset:) Centrosome separation does not occur in a directed manner as detected in the time evolution. (Error bars represent standard deviation)

An improved centrosome separation would require polarized centrosomal movements and can be achieved by a continuous repulsive forces acting between the centrosomes. Evidence suggests that polymerizing microtubules that run into the obstacles, are able to apply pushing force on the obstacles [22, 23, 24]. As the microtubules are uniformly nucleated from the centrosomes, some of the microtubules from one centrosome are capable of exerting an outward force on the opposite centrosome [22, 23, 39] (fig. 3a). In simulation, this pushing (i.e. microtubule-centrosome interaction force or MIF) is generated spontaneously along the line joining the centers of the two centrosomes (fig. 3a). For simplicity, MIF remains constant before it vanishes completely when visibility of centrosomes with respect to each other is blocked by the curvature of the nucleus. Combined effect of MIF and CPF(−) significantly improves the centrosome separation (fig. 3b) in comparison with the sole presence of CPF(−) (fig. 2c). Once MIF stops acting on the centrosomes (corresponding ~100^0^ separation), further separation is driven by CPF(−) which finally brings the centrosomes ~140^0^ apart. Note that magnitude of the separation between the centrosomes at the end of prophase is sufficient to form a stable spindle in the successive stages of the mitosis [40]. Fig. 3c shows trajectories of the centrosomes moving under the effects of MIF and CPF(−). Notice that, initially centrosomes are repealed by each other due to strong MIF dominating over the CPF(−). At larger separation, buckling force 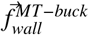 suppresses MIF and CPF(−) takes over; thus give rise to the nonmonotonic trajectories while maintaining the perinuclear positioning of the centrosomes. In simulation, we also check whether MIF alone can separate the centrosomes to the required extent. Fig. 3d shows the centrosomal trajectories for this case, which points out that angular centrosome separation is not initiated and centrosomes lose perinuclear positioning. Thus simulation summarizes that CPF(−) and MIF are jointly responsible for the initial centrosome separation during prophase.

**Figure 3:**
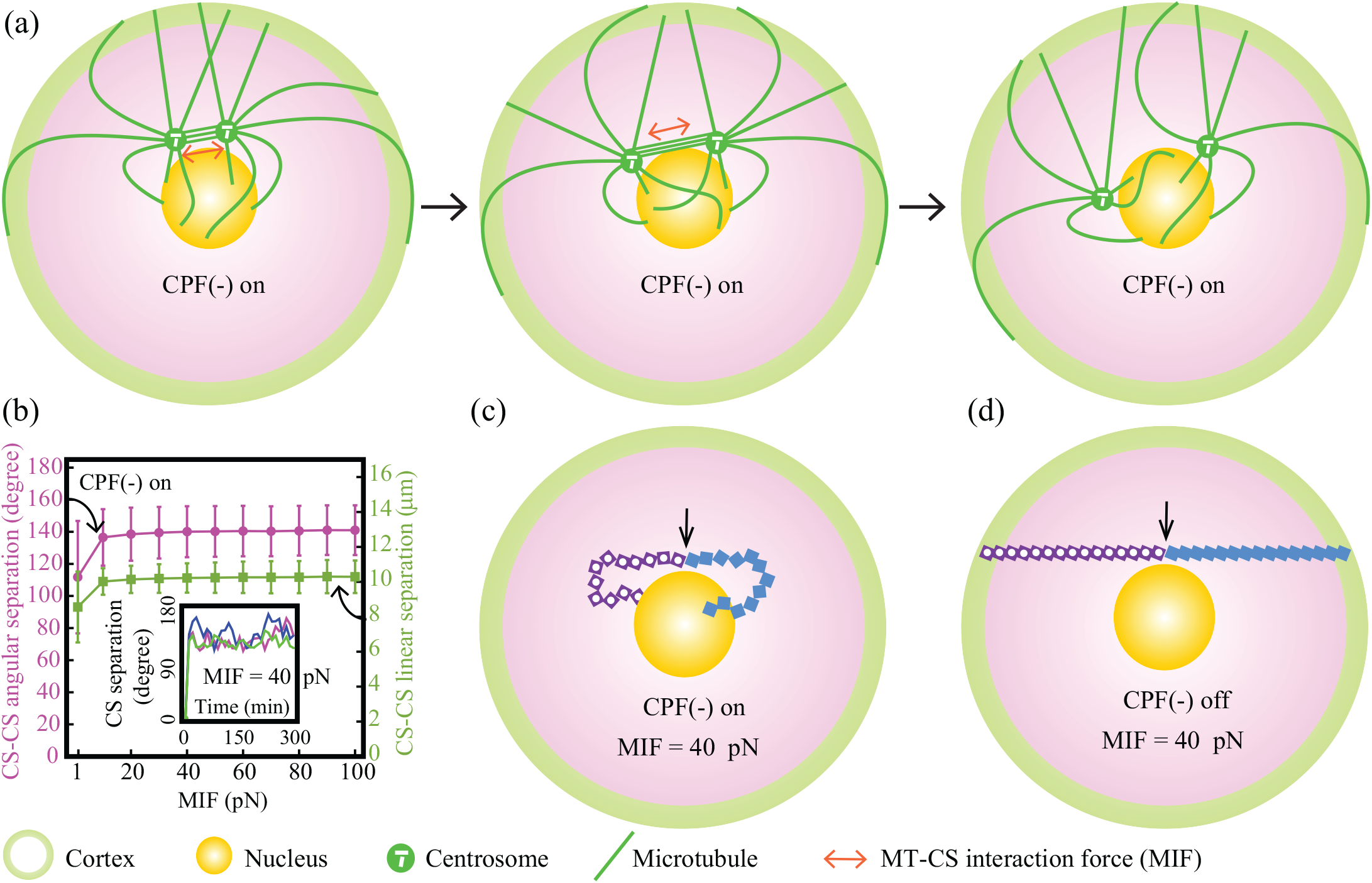
Microtubule-centrosome interaction force (MIF) induces polarity to the centrosomal movements driven by CPF(−). (a) Schematic showing enhanced centrosome separation upon the activation of MIF with CPF(−) on the centrosomes. (b) Quantification of centrosome separation driven by CPF(−) and MIF. (Inset:) Time evolution of three different centrosome separation. In each case, centrosome separation occurs in a directed manner and finally oscillates around the steady state value (~140^0^). (c) and (d) Centrosomal trajectories are plotted upon following the same conventions as they are in fig. 2b. (c) MIF monotonically separates the centrosomes until suppressed by CPF(−) producing nonmonotonic separation. (d) In the absence of CPF(−), only linear centrosome separation occurs due to MIF. (Error bars represent standard deviation)

The key findings of the simulation are: (1) CPF(−) promote the angular movements of the centrosomes around the nucleus and restrict these movements in the perinuclear region, (2) MIF polarizes the centrosomal movements driven by CPF(−). Applying these forces simultaneously, prophase centrosome separation is established. Since a number of evidence supports kinesin-5-independent centrosome separation during prometaphase [41, 42], we further investigate whether it is possible to achieve complete centrosome separation (~180^0^) using CPF(−), MIF coupled with forces to be identified. Evidently, in the state of 180^0^ separation, each centrosome must lie on the antipode of the other centrosome, determined with respect to the nucleus.

Therefore it would be interesting to see the evolution of the centrosome trajectory when an active independent force (AIF) is instantaneously applied on a centrosome toward the antipode of the other. In simulation, we first apply AIF with CPF(−) on the centrosomes and summarize the results in the fig. 4. Fig. 4a is the schematic representation of the centrosome separation upon the activation of AIF with CPF(−) applied on the centrosomes. A complete centrosome separation, plotted in fig. 4b, is reached for AIF >= ~20 pN. The AIF constantly works for centrosome separation and maintains the final separation (fig. 4a). Note that, large AIF (> ~20 pN) suppresses CPF(−), thereby forces the centrosomes to fall on the NE. Thus, final linear separation between the centrosomes decreases marginally and gradually saturates to ~7 µ*m* across the nuclear diameter (fig. 4b). Fig. 4c shows trajectories of the centrosomes gradually separate around the nucleus and finally maintain the separation under the effects of AIF and CPF(−). In our study, it is also observed that (i) AIF alone or (ii) the combination of MIF and AIF or (iii) the combination of CPF(−), MIF and AIF are capable of separating the centrosomes completely. However, in the first two scenarios (i) and (ii), the centrosomes lose the perinuclear positioning and are separated in contact with the NE (data not shown). Thus complete and proper centrosome separation is possible under the combined effect of CPF(−)+AIF or CPF(−)+MIF+AIF.

**Figure 4:**
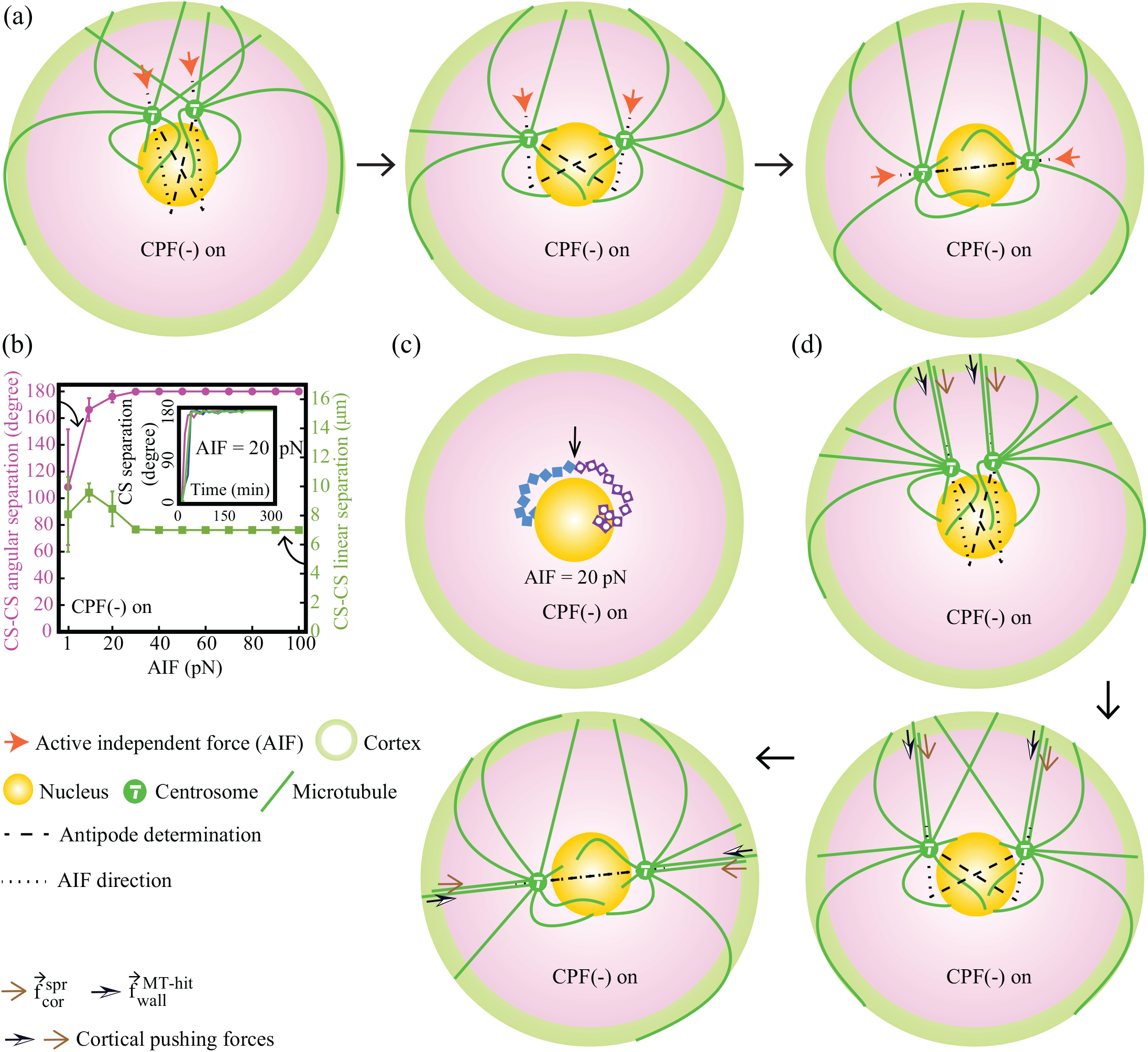
Complete centrosome separation is possible when CPF(−) are accompanied by an active independent force (AIF) pushing each centrosome toward the antipode of the other. (a) Schematic showing complete centrosome separation upon the activation of AIF with CPF(−) applied on the centrosomes. Antipode of individual centrosome is shown by the dashed line and the direction of AIF on each centrosome is shown by the dotted line. (b) Quantification of centrosome separation driven by CPF(−) and AIF. (Inset:) Time evolution of centrosome separation is plotted for three different cases. In each case, centrosome separation occurs monotonically and finally saturates to ~180^0^. (c) Centrosomal trajectories are plotted upon following the same conventions as they are in fig. 3c. In presence of CPF(−) and AIF, centrosomes maintain the perinuclear positioning and are separated completely. (d) Polymerizing microtubules slide along the cell-periphery, and simultaneously cortical pushing forces generate AIF to establish the complete separation. (Error bars represent standard deviation)

Inside the cell, the AIF can be generated through several molecular pathways. Considering the properties of the microtubules, one plausible pathway can be derived from the microtubules polymerizing against the cortex generating active push-ing forces (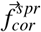 and 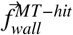) on the centrosome along the length of the microtubules (fig. 4d). Once AIF is initiated, the polymerizing microtubules are able to slide along the cortex producing AIF at successive time steps when centrosomes have repositioned (fig. 4d). This process, iteratively leads to complete centrosome separation and stability of the final configuration. Note that microtubule-sliding along cortex can be initiated either stochastically [33, 43, 24] or by the chemical and physical cues that are induced in the cell from the outer substrate during G2 to M-phase transition [44]. It is identified that a concerted interplay between the extrinsic cues, actin meshwork and the Myo10 motors slides the astral microtubules along the cell-periphery displacing the spindle poles in tandem such that the bipolarity of the mitotic spindle is established [44].

Recently, in the clinical trials for cancer treatments, inhibition of kinesin-5 motors do not show effective success to kill the cancer affected cells [6, 45, 46, 47, 48, 49]. In addition, under survival pressure, the lower eukaryotic cells can have more than one pathway to occur adequate spindle pole separation [39]. From all these outcomes, the respective researchers have now insight that cell generates chemical signal to find out and strengthen the alternative pathway for centrosome separation when one pathway does not function [39, 6]. This chemical signaling is likely to take over the responsibility to slide the polymerizing microtubules along the cell-periphery for supplying AIF at successive time steps after one time generation of the AIF on the centrosomes til an adequate separation is established between them. Future experiments need to be carried out on the AIF in the research field of oncology.

## 4. Discussion

In this paper, we focus on the centrosome separation in mammalian cells during prophase. We show that the combined effect of CPF(−) and MIF facilitates directed movement and lead to a stable separation (~140^0^) of the centrosomes. CPF(−), responsible for the angular movements of the centrosomes around the nucleus, is generated via microtubule-cortex and microtubule-nucleus interactions and MIF, essential for the polarized movements of the centrosomes, is generated via microtubule-centrosome encounter, without requiring kinesin-5 acting between the inter-centrosomal anti-parallel microtubules. Therefore, this study sheds light on the centrosome separation pathways guided by independent centrosomal movement that becomes polarized [4] and achieving stable centrosome separation during prophase without requiring kinesin-5 and dynein [3, 8, 9]. Since the combination of CPF(−) and MIF successfully reproduces prophase centrosome separation, we further investigate whether complete separation can be reached using CPF(−), MIF together with additional active forces. Simulation results indicate that complete separation is possible when CPF(−) work with AIF pushing each centrosome toward the antipode of the other.

For simplicity, our current model does not consider the single molecule dynein interaction with the microtubules [50], instead a mean field approach (average dynein density on the microtubules) [31] is considered. Throughout the simulation, nucleus remains static and concentric with the cell while interacting with the microtubules. However, the position of the nucleus may be off-centered and dynamic due to active forces. A refined model, including these features would be useful. Additionally, implementing deformable cellular objects (nucleus and cell wall) the current model can be made more realistic.

## Acknowledgments

I was supported by fellowship from INSPIRE (IF131156) program from Department of Science and Technology (DST), India and Indian Association for the Cultivation of Science (IACS), Kolkata, India. I gratefully acknowledge Prof. Raja Paul (IACS, Kolkata) for his important comments and inputs to this study.

His Grant No. EMR/2017/001346 of SERB, India and IACS, Kolkata supported the computational facility.

